## Supplemental Figures S1-4, and Table S1 for "The role of mutational spectrum in the selection against mutator alleles"

**Supplementary Information**

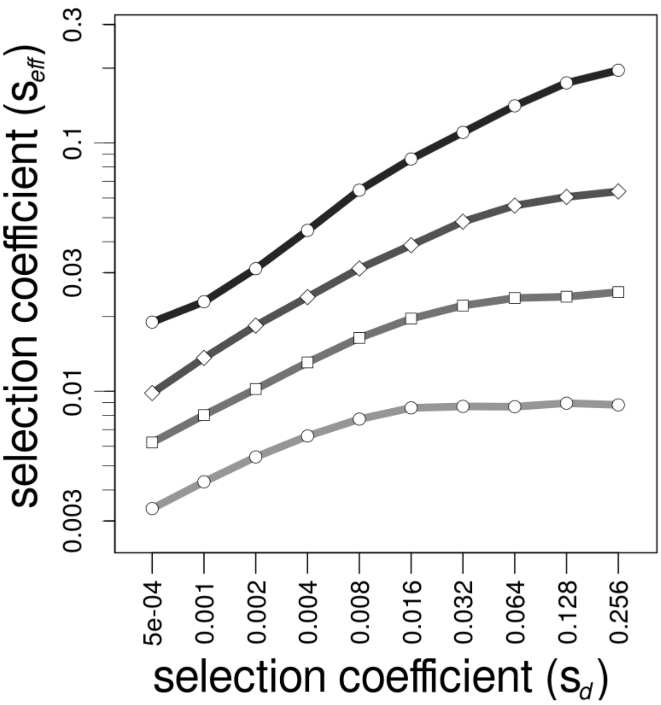


**Fig. S1.-** Dependence of antimutator dynamics on the fitness cost of deleterious mutations. Points represent the effective selection coefficient ($s_{eff}$) of invading antimutator alleles, averaged from 200 independent simulations. Lines depict different values of the mutation rate of the resident mutator (from top to bottom, *m = 1000*; b, *m = 300*; c, *m = 100*; d, *m = 30*). In line with the the Haldane-Muller principle, note how slopes become flatter as mutation rates are the smallest and fitness costs the highest. Other parameters as described in Material and Methods.

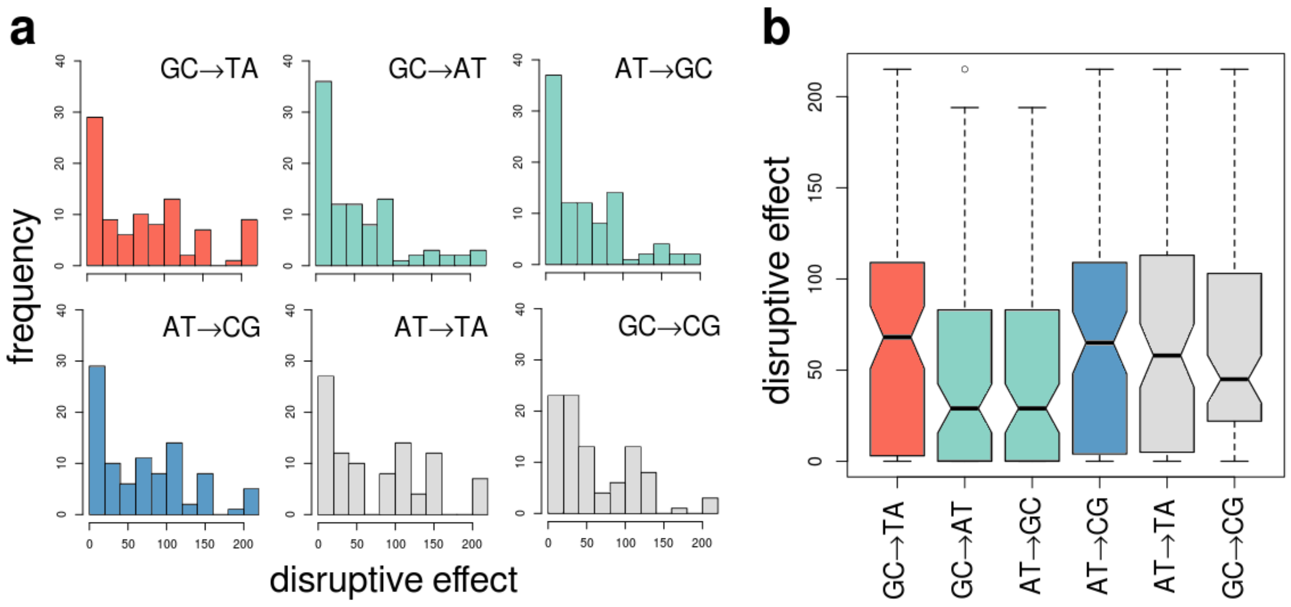


**Fig. S2.-** Protein-disrupting effects of mutations caused by all 6 possible base-substitution mutations across all codons (excluding stop codons). As in Figure 4, colours correspond to the specific mutations that match the mutational spectrum of *mutY*⁻ (red), *mutT*⁻ (blue) and Mismatch Repair⁻ (green) mutators. (a) Histograms showing the distribution of Grantham scores (b) Boxplots of the same scores. Note that, despite average differences, the distribution of scores display a large degree of overlap.

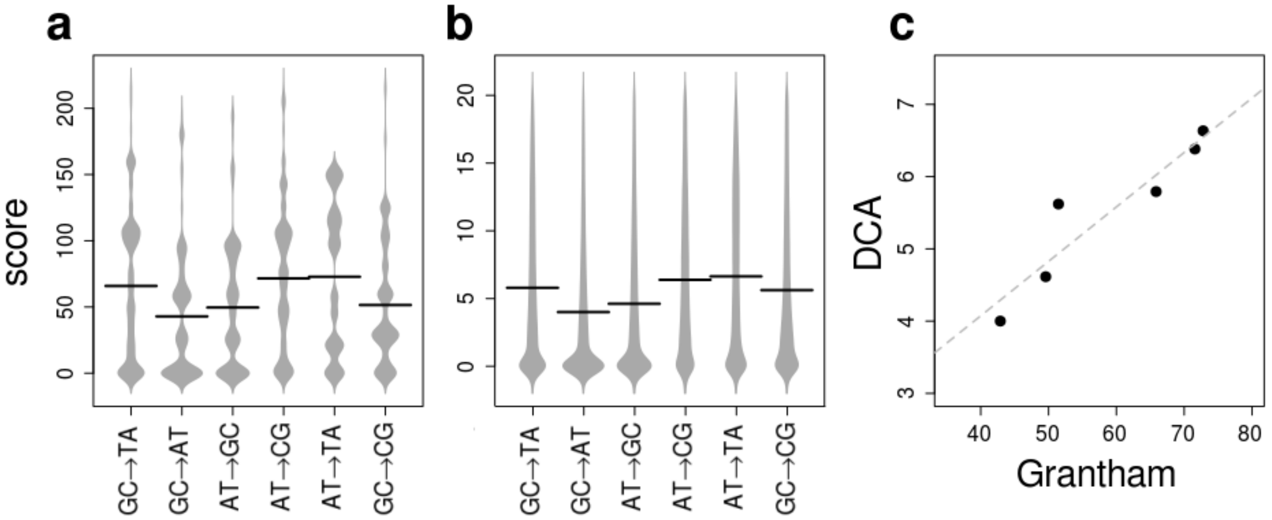


**Fig. S3.-** Comparison of Grantham's matrix with Direct Coupling Analysis (DCA) in predicting the fitness effects of non-synonymous mutations. Data comes from Couce *et al.* (2017), in which both approaches were applied to genomes from the Lenski's Long-Term Evolution Experiment. Only deleterious mutations mapping to empirically validated essential or nearly-essential genes are included here. Panels (a) and (b) show the distribution of scores for Grantham's and DCA respectively. Panel (c) shows the correlation between both methods (*r =0.94*). Plots were drawn using the R library "beanplot" (Kampstra, 2008).

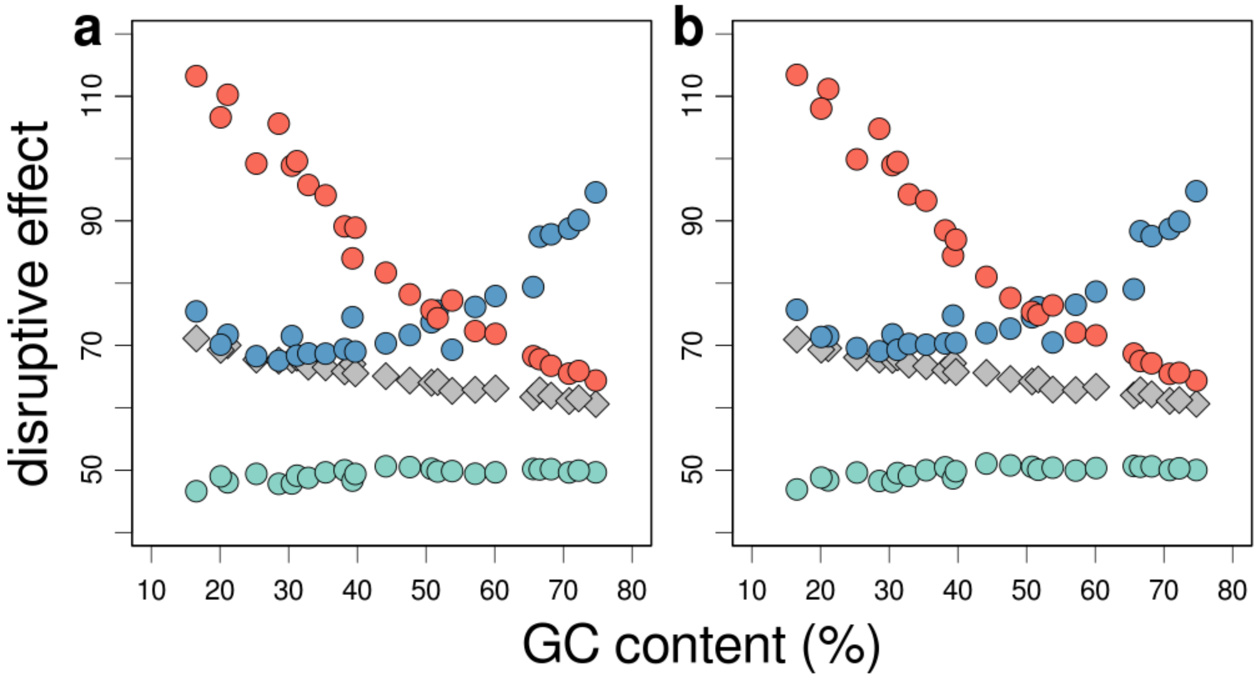


**Fig. S4.-** Average Grantham scores across a panel of species with genomes spanning a wide range of GC compositions. Panels correspond to analyses for genes belonging to the COG categories most commonly enriched in essential genes (H, J and M) (Mandal *et al.*, 2017) (a), and analyses for whole genomes (b). Colours and all other conventions are as in Figure 4.

**Table S1.- Bacterial genomes analyzed in this study**

| **Species Name**[1] | **Gram** | **Taxonomy** | **%GC** | **Size (bp)** | **CDSs** |
|---|---|---|---|---|---|
| *Candidatus Carsonella ruddii PV* | - | γ-proteobacteria | 16.56 | 159662 | 184 |
| *Buchnera aphidicola Cc (Cinara cedri)* | - | γ-proteobacteria | 20.1 | 422434 | 378 |
| *Candidatus Sulcia muelleri CARI* | - | Bacteroidetes | 21.13 | 276511 | 266 |
| *Buchnera aphidicola Bp* | - | γ-proteobacteria | 25.3 | 615980 | 541 |
| *Clostridium difficile CD196* | + | Firmicutes | 28.56 | 4110554 | 3805 |
| *Campylobacter jejuni 4031* | - | є-proteobacteria | 30.47 | 1669329 | 1711 |
| *Prochlorococcus marinus MIT 9312* | - | Cyanobacteria | 31.21 | 1709204 | 2027 |
| *Staphylococcus aureus subsp. aureus NCTC 8325* | + | Firmicutes | 32.87 | 2821361 | 2974 |
| *Bacillus anthracis Ames* | + | Firmicutes | 35.38 | 5227293 | 5844 |
| *Haemophilus influenzae Rd KW20* | - | γ-proteobacteria | 38.15 | 1830138 | 1821 |
| *Helicobacter pylori 2017* | - | є-proteobacteria | 39.3 | 1548238 | 1682 |
| *Streptococcus pneumoniae D39* | + | Firmicutes | 39.71 | 2046115 | 2217 |
| *Xenorhabdus nematophila ATCC19061* | - | γ-proteobacteria | 44.15 | 4587917 | 4770 |
| *Yersinia pestis CO92* | - | γ-proteobacteria | 47.64 | 4829855 | 4587 |
| *Escherichia coli K12* | - | γ-proteobacteria | 50.79 | 4639675 | 4306 |
| *Neisseria meningitidis 053442* | - | β-proteobacteria | 51.7 | 2153416 | 2034 |
| *Corynebacterium glutamicum ATCC 13032* | + | Actinobacteria | 53.81 | 3309401 | 3223 |
| *Brucella abortus A13334* | - | α-proteobacteria | 57.22 | 3286032 | 3566 |
| *Bifidobacterium longum NCC2705* | + | Actinobacteria | 60.12 | 2260266 | 2049 |
| *Mycobacterium tuberculosis Beijing/NITR203* | + | Actinobacteria | 65.61 | 4411128 | 4699 |
| *Pseudomonas aeruginosa PAO1* | - | γ-proteobacteria | 66.56 | 6264404 | 5835 |
| *Burkholderia pseudomallei 1106b* | - | β-proteobacteria | 68.21 | 7224634 | 6990 |
| *Nocardia farcinica IFM 10152* | + | Actinobacteria | 70.83 | 6292344 | 6283 |
| *Streptomyces griseus subsp. griseus NBRC 13350* | + | Actinobacteria | 72.23 | 8545929 | 7547 |
| *Anaeromyxobacter dehalogenans 2CP-1* | - | δ-proteobacteria | 74.72 | 5029329 | 4690 |

[1] as retrieved from Genoscope (*www.genoscope.cns.fr*)
